# A seasonal copepod ‘lipid pump’ promotes carbon sequestration in the deep North Atlantic

**DOI:** 10.1101/021279

**Authors:** Sigrún H. Jónasdóttir, André W. Visser, Katherine Richardson, Michael R. Heath

## Abstract

Estimates of carbon flux to the deep oceans are essential for our understanding for global carbon budgets. We identify an important mechanism, the lipid pump, that has been unrecorded in previous estimates. The seasonal lipid pump is highly efficient in sequestering carbon in the deep ocean. It involves the vertical transport and respiration of carbon rich compounds (lipids) by hibernating zooplankton. Estimates for one species, the copepod *Calanus finmarchicus* overwintering in the North Atlantic, sequester around the same amount of carbon as does the flux of detrital material that is usually thought of as the main component of the biological pump. The efficiency of the lipid pump derives from a near complete decoupling between nutrient and carbon cycling and directly transports carbon through the meso-pelagic with very little attenuation to below the permanent thermocline. Consequently the seasonal transport of lipids by migrating zooplankton is overlooked in estimates of deep ocean carbon sequestration by the biological pump.

## Significance statement

The deep oceans are an important sequestration site for anthropogenic carbon. Every winter, across the North Atlantic, huge numbers of lipid rich zooplankton migrate to the deep ocean (>1000 m) to hibernate. In their migration to overwintering depths, these copepods actively transport lipid carbon and respire it at a rate comparable to the carbon delivered by sinking detritus. This lipid pump is an as yet unresolved component of the sequestration flux of carbon in the oceans, being un-recorded in either direct flux measurements, or estimates based on new production. Unlike other components of the biological pump, the lipid pump does not strip the surface ocean of nutrients, and decouples carbon sequestration from limiting constraints of surface productivity; the lipid shunt.

## Introduction

Understanding the dynamics of deep ocean carbon sequestration is fundamental in estimating global carbon budgets. Atmospheric CO_2_ is taken up by the oceans and some is subsequently fixed as organic carbon by primary producers. The fixed carbon is removed from the surface by several processes described as the biological pump (1) that mainly cover passive sinking of organic detritus. This is an important process that is estimated to sequester 2 - 8 gC m^−2^ yr^−1^ at around 1000m depth (2). Zooplankton are important players in the biological pump, feeding on the primary production which they repackage into fast sinking faecal pellets (3). Other zooplankton mediated processes include the feeding and disruption of particle fluxes (4), and respiration in their daily vertical migrations (5). Here we identify another important role of copepods that plays out mainly in polar regions; the active vertical transport of lipid carbon related to their annual migrations to the deep ocean.

Every year at the end of summer, trillions of copepods descend to the deep ocean basins of the North Atlantic to overwinter in state of diapause (hibernation). Copepods of the genus *Calanus* provide a particularly striking example of organisms exhibiting this life history strategy (6). These species (*C. finmarchicus*, *C. helgolandicus*, *C. glacialis*, *C. hyperboreus* in the polar and temperate North Atlantic and equivalent species in the Pacific and Southern Ocean (7, 8)) form a vital trophic link between primary producers and higher trophic levels (9, 10). In terms of distribution, *C. finmarchicus* is the most cosmopolitan of these and is found in high abundances from the Gulf of Maine to north of Norway (11). Depending on region, the diapause duration of *C. finmarchicus* varies from 4 to 9 months. To survive these long periods, remain neutrally buoyant (12) and fuel their reproduction the following spring (13), these copepods store lipids (up to 70% of their dry weight) in the form of wax esters (WE); long-chain carbon and energy rich compounds that include omega-3 fatty acids. During the diapause period, the animals live at depths from 600 to 1400 m, and temperatures from -1^°^C to 8^°^C (14). Diapause depth varies between regions but it must *a priori* be below the depth of the permanent thermocline to prevent the torpid animals from being prematurely returned to the surface waters.

This manuscript describes a previously unaccounted for mechanism transporting substantial amounts of carbon to the deep North Atlantic by seasonally migrating zooplankton, thus providing an additional explanation of the removal of carbon from the surface oceans. We introduce two new concepts; the lipid pump and the lipid shunt, both of which have important implications for our understanding of ocean carbon cycling.

## Results

Winter surveys (14, 15) have revealed prodigious numbers (15,000 to 40,000 individuals per m^2^; Table 1) of diapause stages copepodite 5 (CV) of *C. finmarchicus* in the various basins across the North Atlantic. While each individual contain only about 200 µg of lipids, the numbers and spatial extent produce a surprisingly large integrated effect presented in Figure 1. In Table 1 we estimate this aspect per basin while in Figure 1 we represent the detailed pattern of the lipid pump, based on individual station estimates as shown in panel c.

**Figure 1.**
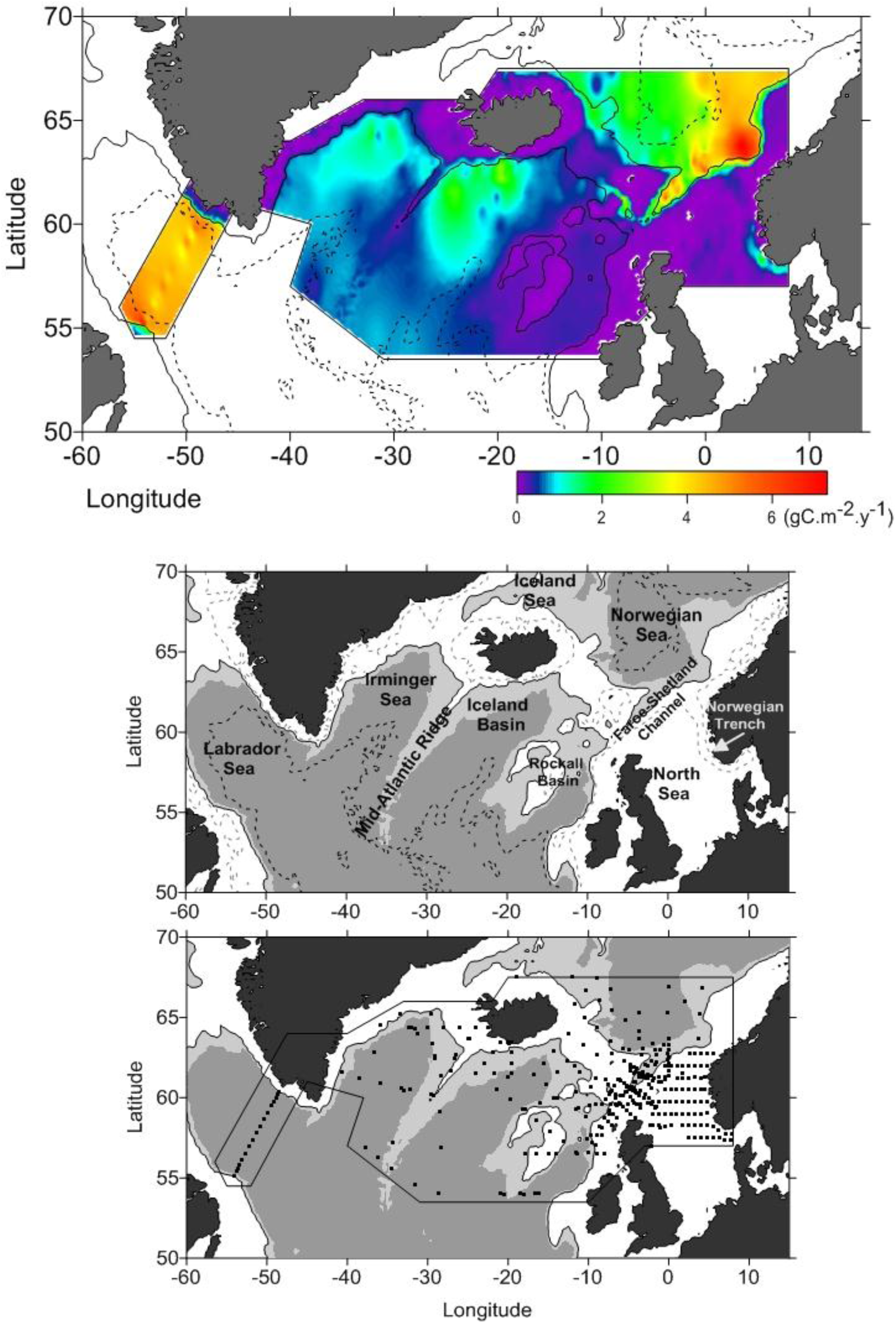
Lipid sequestrated carbon flux. Top panel: A map of carbon flux (gC m^−2^ yr^−1^) associated with lipid sequestration (respiration) of overwintering *Calanus finmarchicus* in the North Atlantic. The carbon flux “hot spots” in the Eastern Norwegian Sea is due to the high copepod abundance but in the Labrador Sea due to larger copepod size and higher respiration due to higher temperatures. The mid panel: Names of the ocean basins referred to in Table 1. Bottom panel: Sampling locations of *Calanus finmarchicus* during winter for abundance, length, overwintering depths and temperatures.

We estimate an annual flux of 1.6 to 6.4 gC m^−2^ yr^−1^ in the form of lipids, actively transported to depth by the annual *C. finmarchicus* migration (Figure 1; Table 1).

**Table 1.**
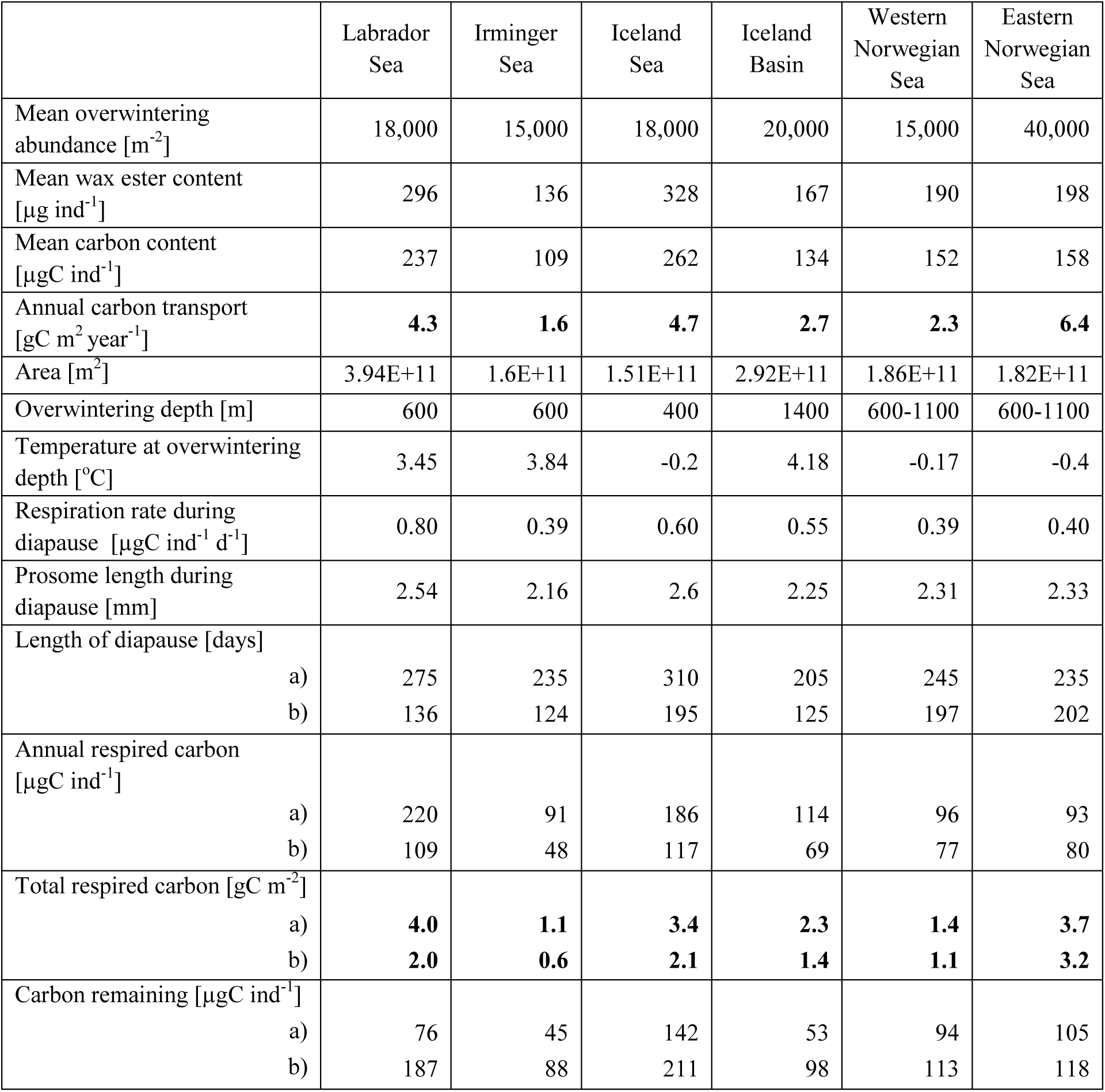
Regional breakdown of the lipid pump calculations. Abundance, overwintering depth, temperatures (14–16). Prosome lengths for the Labrador Sea are published values (16) while the other length measurements are the authors’ own unpublished data. Area of the different ocean basins was estimated using MathWorks mapping tool box. Wax ester content is estimated from the length measurements (17), and for length of diapause we present 2 different estimates a (16) and b (18). See Methods for details. The respective annual respiration for the 2 different models is based on published respiration regressions (17, 19). Carbon remaining is the carbon not utilized over the overwintering period and can be brought back to the surface during spring ascent.

|  | Labrador Sea | Irminger Sea | Iceland Sea | Iceland Basin | Western Norwegian Sea | Eastern Norwegian Sea |
| --- | --- | --- | --- | --- | --- | --- |
| Mean overwintering abundance [m<sup>-2</sup>] | 18,000 | 15,000 | 18,000 | 20,000 | 15,000 | 40,000 |
| Mean wax ester content [µg ind<sup>-1</sup>] | 296 | 136 | 328 | 167 | 190 | 198 |
| Mean carbon content [µgC ind<sup>-1</sup>] | 237 | 109 | 262 | 134 | 152 | 158 |
| Annual carbon transport [gC m<sup>2</sup> year<sup>-1</sup>] | **4.3** | **1.6** | **4.7** | **2.7** | **2.3** | **6.4** |
| Area [m<sup>2</sup>] | 3.94E+11 | 1.6E+11 | 1.51E+11 | 2.92E+11 | 1.86E+11 | 1.82E+11 |
| Overwintering depth [m] | 600 | 600 | 400 | 1400 | 600-1100 | 600-1100 |
| Temperature at overwintering depth [°C] | 3.45 | 3.84 | -0.2 | 4.18 | -0.17 | -0.4 |
| Respiration rate during diapause [µgC ind<sup>-1</sup> d<sup>-1</sup>] | 0.80 | 0.39 | 0.60 | 0.55 | 0.39 | 0.40 |
| Prosome length during diapause [mm] | 2.54 | 2.16 | 2.6 | 2.25 | 2.31 | 2.33 |
| Length of diapause [days] |  |  |  |  |  |  |
| a) | 275 | 235 | 310 | 205 | 245 | 235 |
| b) | 136 | 124 | 195 | 125 | 197 | 202 |
| Annual respired carbon [µgC ind<sup>-1</sup>] |  |  |  |  |  |  |
| a) | 220 | 91 | 186 | 114 | 96 | 93 |
| b) | 109 | 48 | 117 | 69 | 77 | 80 |
| Total respired carbon [gC m<sup>-2</sup>] |  |  |  |  |  |  |
| a) | **4.0** | **1.1** | **3.4** | **2.3** | **1.4** | **3.7** |
| b) | **2.0** | **0.6** | **2.1** | **1.4** | **1.1** | **3.2** |
| Carbon remaining [µgC ind<sup>-1</sup>] |  |  |  |  |  |  |
| a) | 76 | 45 | 142 | 53 | 94 | 105 |
| b) | 187 | 88 | 211 | 98 | 113 | 118 |

The proportion of this transported carbon which remains at depth (i.e. sequestered) depends on mortality and respiration rates. Overwintering temperatures and lipid content determine how long individual copepods can remain in diapause (17, 20) and their respiration rates (17, 19). Estimated respiration rates of 0.4-0.8 µg C ind^−1^ d^−1^ over diapause intervals of 120-300 days, results in 44-93% of the lipid reserves being respired at depth depending on location (Table 1). The sequestration flux associated with respiration of overwintering *C. finmarchicus* is thus estimated to be 1 to 4 gC m^−2^ yr^−1^. Mortality of overwintering copepods, while potentially important to the survival of the *Calanus* population, will in all likelihood augment the fraction of transported carbon remaining at depth, making our sequestration estimate conservative. A portion of the non-respired carbon returns to the surface the following spring with the ascending population. Based on initial values and our estimated respiration loss, the lipid content of individuals at the end of overwintering is about 100 µg C ind^−1^ (Table 1) which is consistent with actual measurements in spring in the North Atlantic (21).

## Discussion

The oceans are an important repository for carbon, and the North Atlantic appears to be particularly active in atmospheric CO_2_ drawdown through both physical and biological process (2). Estimates of the biological pump for the entire North Atlantic range from 1.0 to 2.7 Gt C yr^−1^, constituting 10% to 25% of the global biological pump (2). The large variations in these estimates stem from the application of different observational methods (satellite mounted optical sensors, nutrient budgets, sediment traps, radio nuclides, models and combinations thereof) in their derivation (22). Each of these methods captures some, but not all, of the various processes involved in transporting biogenic carbon to the deep ocean, and this makes the closure of the oceanic carbon budget notoriously difficult (22).

To compare the magnitude of the potential contribution of the seasonal *Calanus* migration to ocean carbon sequestration with that of the biological pump in the North Atlantic, we need both an estimate of the amount of POC leaving the euphotic zone (termed the export flux) (2, 23, 24) and an estimate of how much of this export production sinks to depths below the permanent thermocline where it can be sequestered (25–27). Of the annual mean primary production, a fraction (5% to 15%) (23, 24) is typically exported out of the euphotic zone (∼ 50-100 m), mainly in the form of sinking particulate organic material (faecal pellets and aggregate detritus). A recent review for the North Atlantic (2) provides a range of estimates 29±10 g C m^−2^ yr^−1^ for the regional export flux. There is considerable attenuation of this export flux as it sinks through the mesopelagic to below the permanent thermocline as particles are consumed, fragmented and remineralized. At depths of 600 to 1400 m, only 10% to 20% of the export flux remains (26, 28); i.e. on the order of 2 to 8 gC m^−2^ yr^−1^. Thus, the lipid pump we identify here in association with the seasonal migration of a single zooplankton species, *C. finmarchicus* is of a similar magnitude (1 - 4 g C m^−2^ y^−1^) as the more traditionally studied biological pump.

Many marine organisms execute diel vertical migrations and these are known to regulate trophic interactions and vertical fluxes of carbon (5, 29), nutrients and oxygen (5, 30). In addition to these relatively shallow daily excursions, a number of zooplankton carry out seasonal vertical migrations which allow them to synchronize their life cycles with the seasonal periodicity of primary production in temperate and boreal latitudes.

An important difference between the lipid pump and the traditional biological pump is that the elemental ratios of nitrogen, phosphorus, silicon and iron to carbon are extremely low or zero in lipids resulting in that the lipid pump is essentially decoupled from other nutrient cycles (Figure 2). When copepods synthesize lipids, organic compounds are stripped of their nitrogen and phosphorus, leaving highly reduced carbon chains with much higher C:N and C:P ratios than in typical organic matter and therefore does not strip the surface ocean of limiting nutrients, and decouples the carbon sink from nutrient replenishment rates. The nutrients associated with this lipid storage synthesis are excreted in surface waters long before the overwintering migration takes place. Thus, respiration based on lipids will produce few waste products other than CO_2_ and H_2_O. This results in a “lipid shunt”, equivalent in effect to the C enrichment of POC which occurs in the so-called ‘microbial shunt’ (31). Production of lipids and their transport to depths occurs without any net consumption of the essential limiting nutrients in the surface ocean.

**Figure 2.**
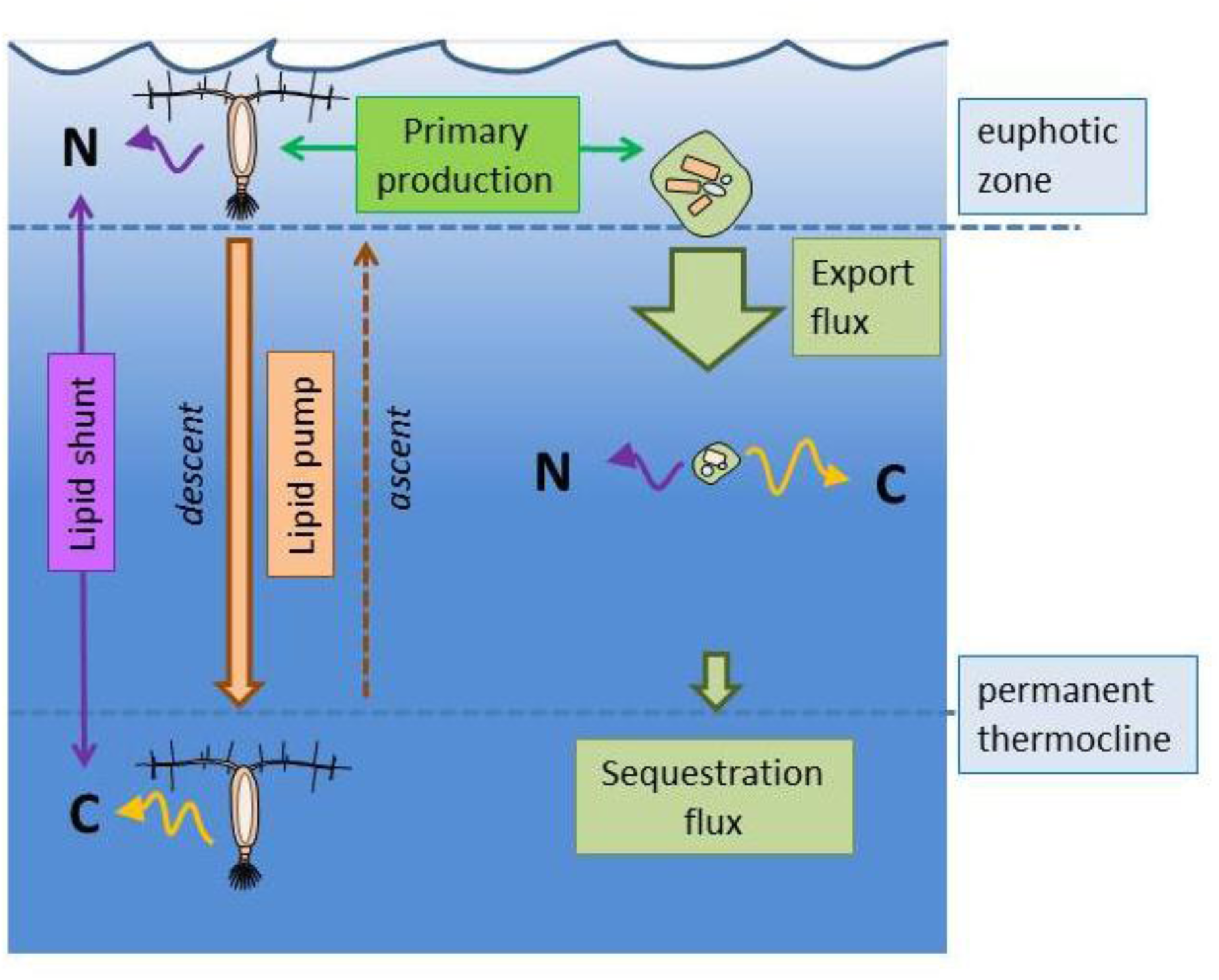
The lipid pump component of the biological pump and the lipid shunt. The export flux of POC is much greater than the active transport of lipids by overwintering copepods. However, much of the POC flux is attenuated as it sinks to depth. At overwintering depths (> permanent pycnocline; 400-1400 m), the lipid transport and the POC flux are comparable (2 to 6 gC m^−2^ yr^−1^). A significant portion of transported lipid is respired at depth (44%-93%). The lipid pump, representing the difference between what descends in the autumn, and what ascends to the surface in spring, is conservatively estimated at (1 to 4 gC m^−2^ yr^−1^), other sources of loss such as predation notwithstanding. The decoupling of nutrient and carbon transport represents a shunt in the biological pump; the nutrients associated with lipid accumulation remain in surface waters, while respired carbon associated with their use over winter, is sequestered at depth.

This study demonstrates that the active vertical transport of lipids by overwintering zooplankton potentially contributes significantly to the ocean’s ability to sequester carbon and act as a carbon sink in the global carbon cycle. The arguments presented here are based on data for a single species and, thus, present a very conservative estimate of the potential importance of this process in carbon cycling. The distribution of *C. finmarchicus* is limited to the North Atlantic. However, ecologically comparable species found in other ocean basins also undertake lipid-fuelled seasonal migrations in their life cycle (8). The potential influence of the lipid pump identified here on global carbon cycling remains to be quantified.

## Materials and methods

Abundances of *C. finmarchicus* in diapause across the N Atlantic, Labrador Sea and Iceland Sea were taken from the original datasets (14, 15). The values from the Iceland Sea are published as dry weight but the absolute numbers were provided by the authors. Body size (prosome length) from overwintering copepods from >300 m depth were obtained from 13 winter cruises in the North Atlantic basins (Figure 1) but published values were used from the Labrador Sea (16). Depth and temperature during diapause is the weighted depth and the corresponding temperature (14).

Storage lipid content of *C. finmarchicus* is taken from the relationship between oil sac volume (OSV) and copepod prosome length (17). We use the mean OSV value for the corresponding size to avoid overestimation in our calculations. We use 0.90 g WE ml^−1^ to convert OSV to individual µg wax esters (32). The carbon weight of WE depends on the fatty acid and alcohol composition, and we base our value on observed relative compositions (33, 34) resulting in 79% of the WE weight being carbon.

Respiration is based on the temperature the copepods experience at depth during diapause and equations established for over-wintering *Calanus* (17). We use the respiration equation that gives µmol O_2_ gC^−1^ h^−1^ and convert it to µgC ind^−1^ by using the length to dry weight relationship for winter and 62% carbon per dry weight (19). Two different model estimates are used to calculate duration of diapause. One is based on respiration and the lipid content (OSV) of the copepods (17) (corrected equation (20)) and is length and temperature depended. The other estimate is based on demographic time series of entry and emergence into diapause from the Labrador Sea, and surface temperatures at the onset of diapause (18).

## Author Contributions

SHJ designed research, SHJ, AWV, MRH performed research; AWV and SHJ wrote the paper with major intellectual input from KR and MRH. All authors discussed the results and commented on the manuscript.

The authors declare no conflict of interest

## Acknowledgement

This work is funded by the Danish Council for Strategic Research under the NAACOS (North Atlantic - Arctic coupling in a changing climate: impacts on ocean circulation, carbon cycling and sea-ice) and EU FP7 program EURO-BASIN; (Basin-scale Analysis, Synthesis & Integration) ENV.2010.2.2.1-1; www.euro-basin.eu).

